# Analysis of the area and the number of pulmonary alveoli through the normal aging process in CD1 mouse

**DOI:** 10.1101/2020.12.11.421032

**Authors:** Marta Ortega-Martínez, Esthefania Gutiérrez-Arenas, Vanessa Gutiérrez-Dávila, Alberto Niderhauser-García, Ricardo M. Cerda-Flores, Gilberto Jaramillo-Rangel

**Affiliations:** Department of Pathology, School of Medicine, Autonomous University of Nuevo Leon, Monterrey, Nuevo Leon, Mexico; School of Biology, Autonomous University of Sinaloa, Culiacan, Sinaloa, Mexico; School of Nursing, Autonomous University of Nuevo Leon, Monterrey, Nuevo Leon, Mexico

## Abstract

During the aging process, the lung exhibits structural changes accompanied by a decline in its function. The related information currently available is still scarce and contradictory. In addition, changes in some pulmonary parameters through aging process are species- and strain-dependent. The aim of this study was the assessment of the area and the number of pulmonary alveoli through the normal aging process in CD1 mouse. Paraffin-embedded sections of lungs from CD1 mice at age of 2, 6, 12, 18, or 24 months were stained with hematoxylin and eosin and examined using a light microscope. Images were captured using a camera linked to an image analysis software to measure areas and count alveoli. There was a significant difference in the alveolar area among the ages analyzed (F=87.53, Sig.=0.000). The alveolar area of the 6-, 12-, 18-, and 24-month-old mice was significantly greater (all *p* values < 0.001) than in mice at 2 months of age. Also, the alveolar number was significantly different among the ages tested (F=3.21, Sig.=0.023). The number of alveoli in mice at 2 months of age was greater than in mice at all other age groups, reaching statistical significance when compared with the 6-, 12-, and 18-month-old mice (*p* values of 0.044, 0.014, and 0.002, respectively). Thus, we observed an increase in alveolar area and a decrease in alveolar number through the aging process. This information might be useful to understand pathologic changes underlying susceptibility of elderly individuals to chronic lower respiratory tract diseases.

## Introduction

Aging process is associated with structural changes in the lung that lead to a decrease in its function. These alterations may increase susceptibility of elderly individuals to chronic lower respiratory tract diseases, such as asthma and chronic obstructive pulmonary disease (COPD) [1,2]. Lung aging-related research has been devoted mainly to the analysis of alveolar structure. There is almost a consensus that the principal feature of the senile lung is the homogeneous enlargement of alveolar airspaces [3–5]. Information available in relation to the number of alveoli through aging is more limited and controversial [6–8].

Interestingly, it has been found that changes in alveolar area and number through aging process are species- and strain- dependent. For example, it has been reported that the human lung maintains a very constant number of alveoli as it ages [6], whereas dogs [9] and female rhesus macaques [8] suffer a decrease in it. In contrast, a study found that senile rats do not show alterations in this parameter [10]. Also, it has been suggested that BALB/cNNia mice may be resistant to an aging-related increase in alveolar size [11], whereas DBA/2 and C57BL/6 mice display an air space enlargement during late life stages [12,13]. These events could be related to innate pulmonary variations between mouse strains; for example, it has been described that C3H/HeJ, C57BL/6J and A/J mice differ naturally in alveolar size [14].

Although the mouse strain CD1 has been widely used to examine multiple aspects of the general aging process [15–17], its alveolar morphometric analysis has not been reported yet. We previously evaluated the cell turnover of the bronchiolar epithelium and analyzed the non-epithelial areas of bronchioles during the aging process in this mouse strain [18,19]. The aim of this study was the assessment of the area and the number of pulmonary alveoli through the normal aging process in CD1 mouse.

## Materials and methods

### Animals and experimental design

Animals and experimental design were as described previously [18,19]. Male CD1 mice were maintained in standard conditions: stainless-steel cages, 12:12 h day–night cycle, 55–60% relative humidity, temperature 18–21°C, and with food and water *ad libitum*. Three animals were instantly sacrificed at the age of 2, 6, 12, 18 or 24 months by cervical dislocation. Only the right lungs were processed and analyzed in order to more consistently control the selection of samples. Animal care was provided according to the principles and procedures defined in the National Research Council Guide for the Care and Use of Laboratory Animals (8th edition), and in the Mexican Guidelines ZOO-062. The protocol was approved by the Institutional Review Board and Ethics Committee of the School of Medicine of the Autonomous University of Nuevo Leon (No. PA19-00001).

Lungs were fixed in 10% neutral buffered formalin and embedded in paraffin. Five micrometers serial sections were cut, deparaffinized in xylene and hydrated in a graded series of alcohol. Each section was stained with hematoxylin and eosin (H&E) according to standard techniques. Three tissue sections per animal taken from the middle of the lung were used, so that a portion of each lung lobe was included in each section for its analysis.

### Assessment of area and number of alveoli

H&E sections were used for estimation of alveolar area and number. Each section was divided into four equal quadrants and a field of microscopy per quadrant was randomly considered, evaluating only the alveoli and discarding large airways and blood vessels. Only closed alveoli were evaluated; alveoli that opened to an alveolar duct were not considered (Fig 1). Sections were examined using a Primo Star light microscope (Carl Zeiss Microscopy GmbH, Oberkochen, Germany), and high-resolution color images (x400) were captured using a Axio-Cam ICc1 camera (Carl Zeiss Microscopy GmbH) linked to image analysis software Zen lite 2011 (Carl Zeiss Microscopy GmbH) to measure areas and count alveoli. Identifying information on each slide was temporarily masked, until after morphometric analysis by a single observer was completed.

**Fig 1.**
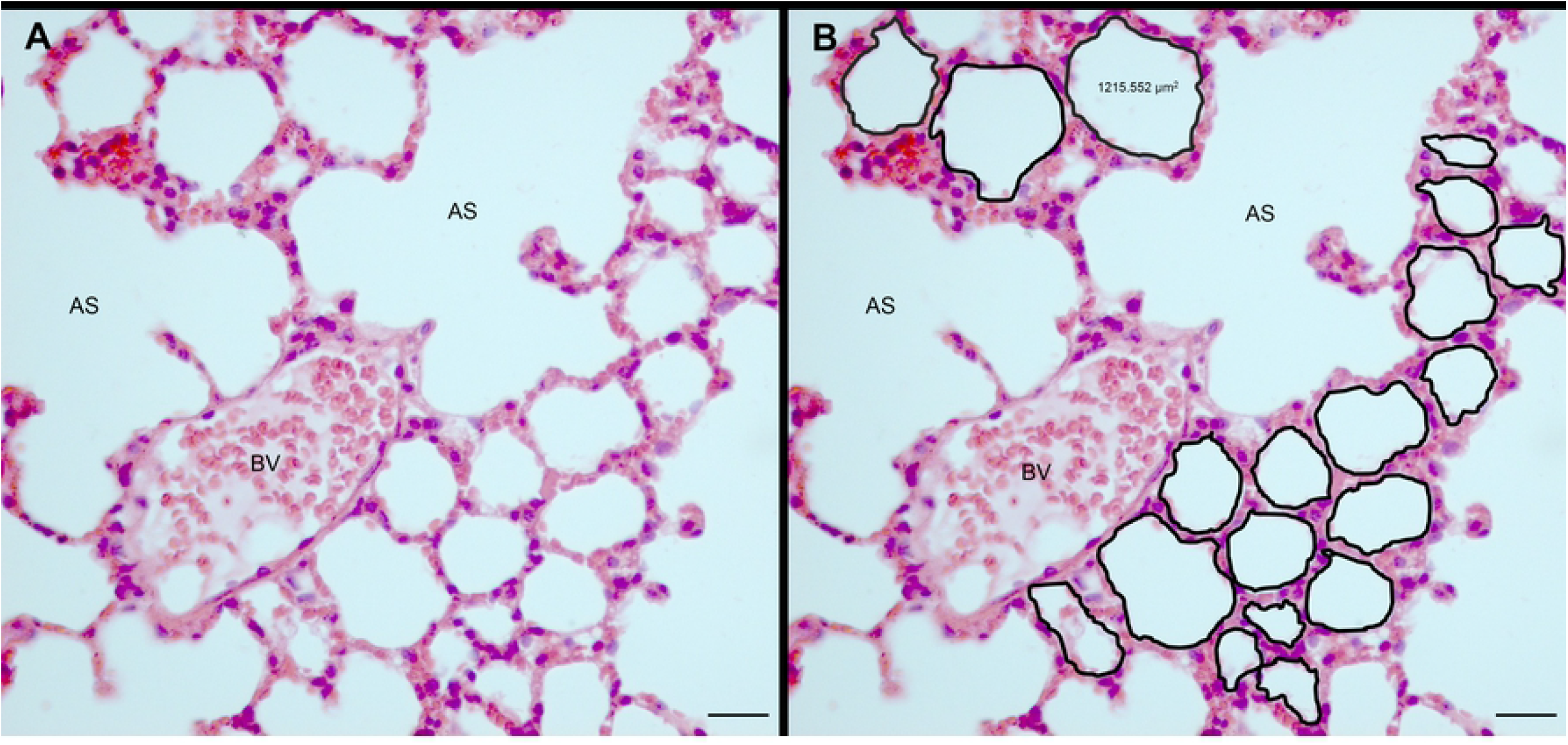
Analysis of alveolar area and number from healthy 2-,6-, 12-, 18-, and 24-month-old CD1 mice. (A) Representative lung tissue section from CD1 mice stained with hematoxylin and eosin. (B) The area measured was the surface enclosed by a black line, as shown in the alveoli (see example in the upper middle part of the figure). All alveoli present in the microscopic field were considered for the count (there are 19 alveoli in this example). AS, alveolar sac; BV, blood vessel. Bars, 20 μm.

### Statistical analysis

The results are presented as means ± 1 standard error (SE). One-way ANOVA test was used to determine statistical significance (*p* < 0.05), followed by post hoc analysis using the least significant difference (LSD) test when significant differences were found between groups. The data were analyzed using the SPSS for Windows software (SPSS, Inc., Chicago, IL, USA), release 21.0.

## Results

The results are summarized in Table 1. On average, 85 alveoli per slide were measured and counted (range 32-178). The results of the ANOVA test showed a significant difference in the alveolar area (AA) among the ages analyzed (F= 87.53, Sig.= 0.000). LSD test revealed that the AA of the 6-, 12-, 18-, and 24-month-old mice (598±12, 668±18, 881±20, and 743±21 μm^2^, respectively) was significantly greater (all *p* values < 0.001) than in mice at 2 months of age (488±11 μm^2^). The AA in mice at 24 months of age was significantly higher than in the 6- and 12-month-old mice (*p* values < 0.001, and 0.002, respectively). Mice at 18 months of age had a significantly higher AA than mice at 6 and 12 months of age (both *p* values < 0.001). Also, mice at 12 months of age had a significantly higher AA than in mice at 6 months of age (*p* value of 0.002). Finally, the AA in mice at 24 months of age was significantly smaller than in the 18-month-old mice (*p* value < 0.001) (Fig 2). The alveolar number (AN) was significantly different among the ages tested (F= 3.21, Sig.= 0.023). LSD test revealed that the AN of the 6-, 12-, and 18-month-old mice (80.6±3.7, 74.8±4.5, and 65.1±7.3 alveoli, respectively) was significantly smaller (*p* values of 0.044, 0.014, and 0.002, respectively) than in mice at 2 months of age (105.3±14.6) (Fig 3).

**Table 1.** Morphometric parameters of pulmonary alveoli from CD1 mouse through the normal aging process.

| Parameter | Age (months) |  |  |  |  |
| --- | --- | --- | --- | --- | --- |
|  | 2 | 6 | 12 | 18 | 24 |
| Area ($\mu\text{m}^2$) | 488±11 | 598±12<sup>a</sup> | 668±18<sup>a,b</sup> | 881±20<sup>a,b,c</sup> | 743±21<sup>a,b,c,d</sup> |
| Number | 105.3±14.6 | 80.6±3.7<sup>a</sup> | 74.8±4.5<sup>a</sup> | 65.1±7.3<sup>a</sup> | 87.5±6.9 |

Data reported as mean ± standard error (SE). Data were analyzed by a one-way ANOVA followed by a least significant difference test. Results were considered statistically significant at $p < 0.05$ .
<sup>a</sup>Statistically significant compared with 2-month-old mice.
<sup>b</sup>Statistically significant compared with 6-month-old mice.
<sup>c</sup>Statistically significant compared with 12-month-old mice.
<sup>d</sup>Statistically significant compared with 18-month-old mice

**Fig 2.**
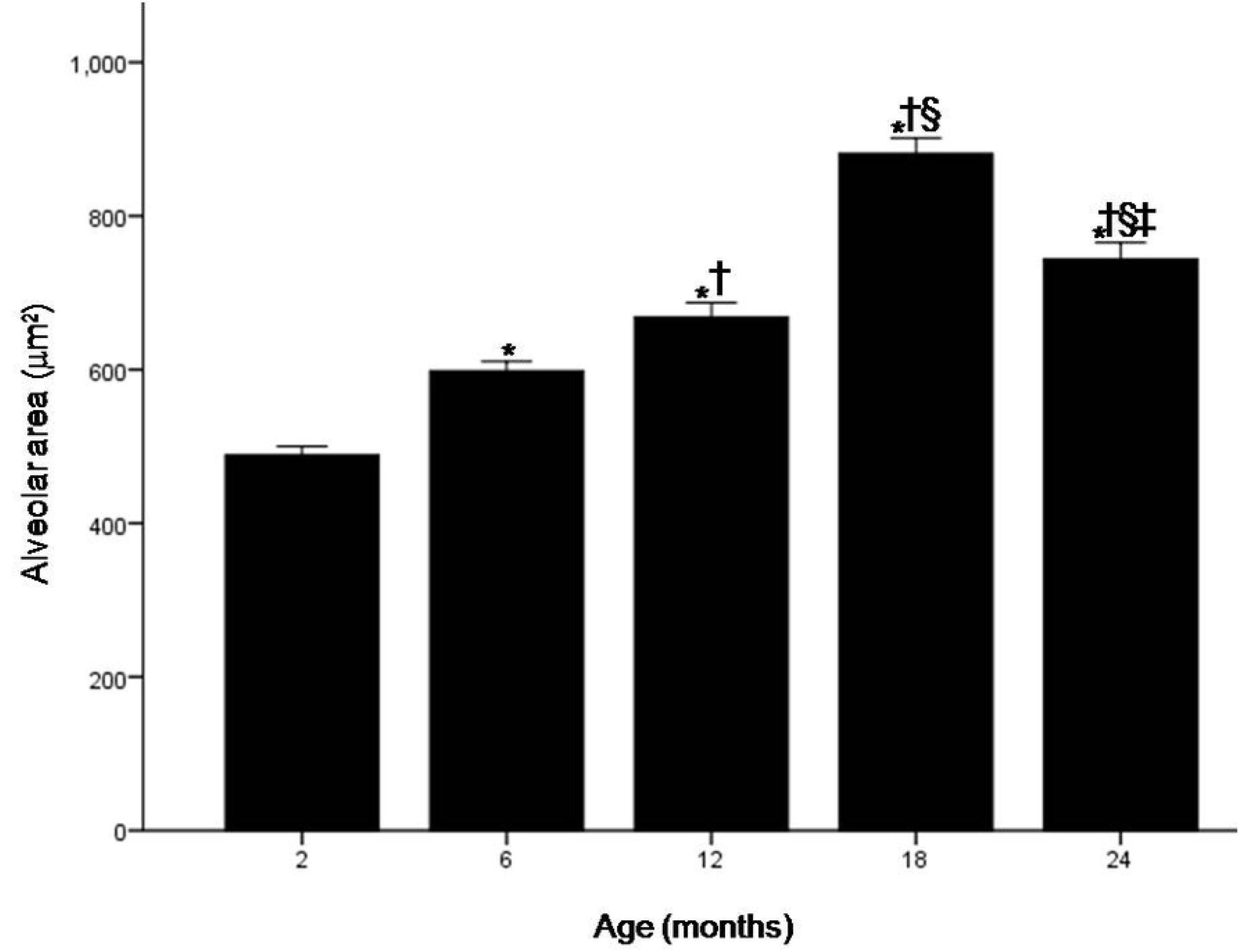
Alveolar area (AA) from healthy 2-,6-, 12-, 18-, and 24-month-old CD1 mice. The AA of the 6-, 12-, 18-, and 24-month-old mice was significantly greater (all *p* values < 0.001) than in mice at 2 months of age (*). The AA in mice at 24 months of age was significantly greater (at most *p* = 0.002) than in the 6- (†), and 12-month-old mice (§). Mice at 18 months of age had a significantly higher AA than mice at 6 (†) and 12 (§) months of age (both *p* values < 0.001). Also, mice at 12 months of age had a significantly higher AA than in mice at 6 months of age (†; *p* = 0.002). Finally, the AA in mice at 24 months of age was significantly smaller than in the 18-month-old mice (‡; *p* < 0.001). Values are expressed as means ± 1 standard error. Data were analyzed by a one-way ANOVA followed by a least significant difference test. Results were considered statistically significant at *p* < 0.05.

**Fig 3.**
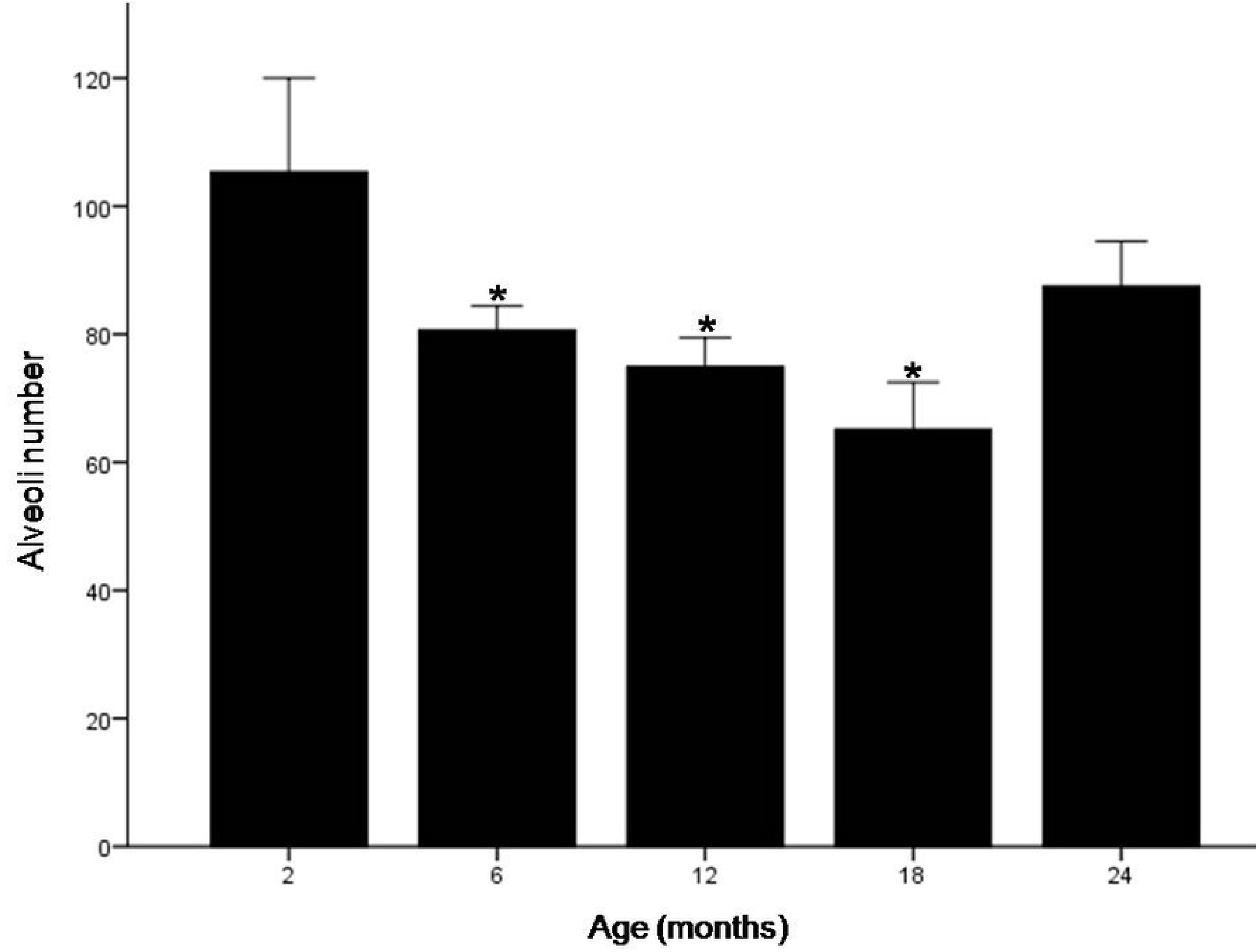
Alveolar number (AN) from healthy 2-,6-, 12-, 18-, and 24-month-old CD1 mice. The AN of the 6-, 12-, and 18-month-old mice was significantly smaller than in mice at 2 months of age (*; at most *p* = 0.044). Values are expressed as means ± 1 standard error. Data were analyzed by a one-way ANOVA followed by a least significant difference test. Results were considered statistically significant at *p* < 0.05.

## Discussion

During the aging process, the lung exhibits structural changes accompanied by a decline in its function. Although alveolar changes have been the most widely studied phenomenon in this area, the information currently available is still scarce and contradictory. In addition, the results obtained vary according to the species and the strain analyzed. The aim of this study was the assessment of the area and the number of pulmonary alveoli through the normal aging process in CD1 mouse.

Our findings revealed that alveolar area increased and number of alveoli decreased with age. Alveolar area increased significantly among all ages between 2 and 18 months (Table 1 and Fig 2), while the 2-month-old mice had a significantly higher number of alveoli when compared to mice at 6, 12, and 18 months of age (Table 1 and Fig 3).

The results obtained in most of the previous studies coincide with ours (Table 2), since in them the lungs of the old animals also presented larger alveoli when compared with the younger ones [3,4,10,12,13,20,21]. The alveolar enlargement observed in the aged lung has been mainly attributed to a loss of elastin fibers in alveolar walls [3,5]. However, since several authors did not observe loss of elastin in the senile lung [4,10] more research is necessary in this area. It is likely that other factors contribute to the enlargement of alveoli through the aging process, such as oxidative stress and cell death [12].

**Table 2.** Changes in size and number of pulmonary alveoli in elderly subjects in comparison with younger ones in different species and strains.

| **Size** |  |  |
| --- | --- | --- |
| **Specie/Strain** | **Result** | **Reference** |
| Fischer-344 rats | Increased | [20] |
| BALB/cNNia mice | No changes | [11] |
| Fischer-344 rats | No changes | [10] |
| Sprague Dawley rats | Increased | [10] |
| Human | Increased | [3] |
| Wistar rats | Increased | [4] |
| Fischer-344 rats | Increased | [21] |
| DBA/2 mice | Increased | [12] |
| C57BL/6 mice | Increased | [13] |
| **Number** |  |  |
| **Specie/Strain** | **Result** | **Reference** |
| Human | No changes | [6] |
| Human | No changes | [7] |
| Beagle dogs | Decreased | [9] |
| F344 rats and Sprague Dawley rats | No changes | [10] |
| Female rhesus macaques | Decreased | [8] |

On the other hand, we observed that the number of alveoli decreased through aging, which is consistent with other reports [Table 2;8,9]. At the moment, there does not exist an explanation for this diminishment in the alveolar number. Using corrosion models, Pump [22] observed an enlargement and fusion of alveoli in elderly men. If such a phenomenon also occurs during the aging process in the mouse lung, this could explain the decrease in the alveolar number accompanied by the alveolar enlargement reported here.

Interestingly, we found a significant decrease in alveolar area between the 18- and 24-month-old mice (Fig 2) and an increase in alveolar number between the same ages, although it was not statistically significant (Fig 3). We and other authors previously reported changes in the lung that were nonlinear through time [12,18–20]. For now, there is no explanation for these events. The analysis of the differential gene expression profiles in lung during aging and the availability of large transcriptomic/proteomic databases might contribute to explain these findings [23].

In contrast to our results, other authors found no difference in alveolar size [10,11] or in the number of alveoli [6,7,10] between aged individuals and young subjects (Table 2). These discrepancies could be related to methodological aspects. We used an image analysis software to asses directly both alveolar area and number in lung transversal cross-sections. This morphometric method might be more exact in comparison with other strategies that obtain results indirectly based on mathematical equations that assume a specific shape of the airspace and require complex calculations. Alternatively, structural and functional pulmonary characteristics that are strongly inherent to the species and strains analyzed (Table 2) might be the cause of the discrepancies. Molecular techniques, as the analysis of the expression of genes involved in the lung aging process in each species or strain, might be helpful to clarify the differences.

In conclusion, here we report, for the first time, the assessment of the area and the number of pulmonary alveoli in a broad range of ages in CD1 mouse. We observed an increase in alveolar area and a decrease in alveolar number through the aging process. This information might be useful to understand pathologic changes underlying susceptibility of elderly individuals to chronic lower respiratory tract diseases. More research is needed to know the molecular mechanisms that give rise to the findings described here.

## Author Contributions

**Conceptualization:** Gilberto Jaramillo-Rangel, Marta Ortega-Martínez

**Data curation:** Marta Ortega-Martínez, Ricardo M. Cerda-Flores

**Formal analysis:** Ricardo M. Cerda-Flores, Vanessa Gutiérrez-Dávila

**Investigation:** Marta Ortega-Martínez, Esthefania Gutiérrez-Arenas

**Methodology:** Marta Ortega-Martínez, Esthefania Gutiérrez-Arenas, Vanessa Gutiérrez-Dávila, Gilberto Jaramillo-Rangel

**Project administration:** Gilberto Jaramillo-Rangel, Alberto Niderhauser-García

**Resources:** Gilberto Jaramillo-Rangel, Alberto Niderhauser-García

**Supervision:** Gilberto Jaramillo-Rangel

**Validation:** Marta Ortega-Martínez, Esthefania Gutiérrez-Arenas, Vanessa Gutiérrez-Dávila, Ricardo M. Cerda-Flores

**Visualization:** Marta Ortega-Martínez, Esthefania Gutiérrez-Arenas, Vanessa Gutiérrez-Dávila

**Writing – original draft preparation:** Marta Ortega-Martínez, Gilberto Jaramillo-Rangel, Esthefania Gutiérrez-Arenas

**Writing – review & editing:** Marta Ortega-Martínez, Gilberto Jaramillo-Rangel, Vanessa Gutiérrez-Dávila

